# Redirecting carbon and electron flow in methanogenic laboratory scale anaerobic digestors with hypophosphite

**DOI:** 10.64898/2026.09.04.749529

**Authors:** Matt E. Weaver, Ruiwen Hu, Ambrose Wang, Yi Liu, John D. Coates, Hans K. Carlson

## Abstract

In anaerobic digestion, formate is a central electron carrier linking primary and secondary fermentation to methanogenesis. Efforts to alter methane production have largely focused on directly targeting methanogens, while disruption of other trophic levels is underexplored. We recently demonstrated that hypophosphite is a naturally occurring inhibitor of formate exchange in syntrophic methanogenic systems. Here, we investigated whether intercepting formate exchange using hypophosphite can modulate methane production in a complex fermentative methanogenic system and what the longer-term consequences are for the structure and function of the microbiome. Using continuous up-flow anaerobic sludge blanket (UASB) columns fed with whey, we found that 100 µM hypophosphite resulted in a transient (17 days) reduction in methane production by 20% to 50%, while carbon and electron flow was redirected towards hydrogen and volatile fatty acids. Accumulation of propionate, butyrate, branched-chain fatty acids, and not acetate indicated disruption of syntrophic formate metabolism. Following recovery of methane production in hypophosphite treated columns, we increased hypophosphite concentrations to millimolar levels and saw no repeated suppression of methane production. 16S rDNA amplicon sequencing revealed that hypophosphite treatment did not cause significant changes to microbiome composition. Methanogenic activity assays revealed a loss of formatotrophic methanogenic capacity in hypophosphite treated columns. Together, these results show that specific inhibition of formate exchange between syntrophs and methanogens decouples fermentative and methanogenic processes without altering microbiome composition with carbon and electron flow to methane ultimately redirected through hydrogen or acetate. This work demonstrates that hypophosphite can intercept syntrophic formate exchange in an engineered system and that long-term treatment with hypophosphite redirects electron flow towards electron carriers other than formate.

## Introduction

Methane (CH₄) is a potent greenhouse gas with a global warming potential approximately 28 times greater than carbon dioxide (CO₂) over a 100-year timescale (Smith et al., 2014). Nearly 70% of the CH4 emitted around the globe is produced by methanogenic archaea, of which over a quarter of the substrates come from anthropogenic sources (Lyu et al., 2018). Developing new mitigation strategies to reduce biogenic CH4 production in these systems and understanding the impact they have on the microbial communities involved is essential in reducing the anthropogenic carbon footprint.

The most common strategy for reducing methane production has been to directly inhibit methanogens by targeting their ability to generate CH₄. A primary target is methyl coenzyme M reductase (MCR), the enzyme that catalyzes the terminal step of methanogenesis. Coenzyme M (CoM) analogs such as 2-bromoethanesulfonate (BES), 2-chloroethanesulfonate, and 3-nitroxypropanol (3-NOP) competitively inhibit MCR at its active site, thereby blocking methane formation (Chidthaisong and Conrad, 2000; Ungerfeld et al., 2004; Siriwongrungson et al., 2007; Hristov et al., 2015; Duin et al., 2016; Jeong et al., 2024). While effective, these approaches tend to be expensive when unsubsidized making them difficult to apply outside of controlled studies (Chae et al., 2010; Pupo et al., 2025). Moreover, they focus exclusively on methanogens and do not address the broader metabolic network that supplies substrates for methane production. Targeting shared metabolic intermediates that link methanogens to methanogen-associated microorganisms offers an alternative systems-level strategy to redirect carbon and electron flow away from methane formation.

Continuous-flow reactors provide a tractable approach for evaluating how novel mitigation strategies influence CH₄-producing microbial consortia in both natural and engineered systems. Up-flow anaerobic sludge blanket (UASB) reactors are among the most common anaerobic continuous-flow reactors and are typically fed mixes of waste (Tawfik et al., 2008; Abdelgadir et al., 2014; Mao et al., 2015). Upon entering the reactor, influent waste contacts a sludge bed of granules composed of extracellular polymeric substances and the microbial consortium required to break down the waste (Díaz et al., 2006; Mills et al., 2021). In the absence of oxygen and other exogenous electron acceptors, the organic waste is degraded to CH4 and CO2. Polymeric waste is first broken down into monomers, so that primary fermentation can then transform the monomeric waste to produce CO2, formate, hydrogen (H2), acetate, alcohols, and (di-)carboxylic acids (Leschine, 1995; Blair et al., 2021; Yu et al., 2023). The mix of alcohols and carboxylic acids are further fermented to methanogenic substrates (i.e., acetate, formate, and H2) by secondary (syntrophic) fermenters, which are thermodynamically dependent on the methanogens consuming their products to generate energy from their reactions (Schink, 1997; McInerney et al., 2009). Due to the movement of carbon and electrons through metabolically coupled reactions to reach CH4, perturbations at any of these trophic levels could cause a cascade of uncoupling that leads to a reduction in CH4 production.

In anoxic environments, formate is a metabolite important in both anabolic and catabolic reactions. Anabolically, it is important for the synthesis of purine nucleotides and production of the central metabolite acetyl-CoA (Knappe and Sawers, 1990; Marolewski et al., 1994; Jordan and Reichard, 1998). Catabolically, formate is generated during primary and secondary fermentation and is subsequently consumed by methanogens as an electron donor for methane production (Bae and McCarty, 1993). Due to this ubiquity across all trophic levels of anaerobic digestion, formate represents a shared metabolic currency and potential target for redistributing carbon and electron flow away from methane at several trophic levels.

A structural analog of formate found in the environment is hypophosphite, a reduced inorganic form of phosphorus (Takamiya, 1953a; McDowell et al., 2004; Morton et al., 2005; Pasek et al., 2014). Early work demonstrated that hypophosphite can disrupt the activity of formate-dependent enzymes and transport systems in *Escherichia coli*, including formate dehydrogenase (FDH), pyruvate–formate lyase (PFL), and the formate transporter FocA (Takamiya, 1953b; Knappe et al., 1984; Brush et al., 1988; Plaga et al., 1988; Unkrig et al., 1989; Suppmann and Sawers, 1994). Other bacterial fermenters grown in pure culture were later shown to be perturbed by hypophosphite at millimolar concentrations (Seward et al., 1982; Wood et al., 1986; Jeffery Rhodehamel and Pierson, 1990; Rhodehamel and Pierson, 1990; Rydzak et al., 2014; Lee et al., 2025). Consistent with these enzyme-level effects, experiments using washed syntrophic–methanogenic cell suspensions suggest that hypophosphite (1 mM) can inhibit formate-exchange but not hydrogen exchange (Sieber et al., 2014). Agne and colleagues showed that hypophosphite can inhibit *Syntrophus aciditrophicus* grown axenically on crotonate by targeting a membrane-bound FDH (Agne et al., 2022). Recent work from our group demonstrated that hypophosphite is a selective inhibitor of syntrophic methanogenic formate exchange in natural anaerobic ecosystems, including rice fields and ruminant systems at low micromolar concentrations (<10µM) (Hu et al., 2026). Physiological assays with a model methanogen further implicated methanogenic formate metabolism as an important target of hypophosphite.

In this study, we combine continuous-flow culturing and 16S rDNA amplicon sequencing to ascertain how hypophosphite disrupts an anaerobic CH4-generating community from an engineered system. We seeded room-temperature UASB columns with sludge from an anaerobic digester and fed them whey, a common byproduct of cheese production (Haast et al., 1985). To monitor disruption, we measured the chemical oxygen demand (COD), gas production, and volatile fatty acid (VFA) composition in hypophosphite treated and untreated columns over 52 days. Sub-millimolar concentrations of hypophosphite were initially sufficient to reduce methane production by ∼30% while driving the accumulation of hydrogen and VFAs prior to adaptation and rebound of methane production. Further increases in hypophosphite concentration produced no subsequent measurable effect on methane production. 16S rDNA amplicon sequencing revealed no major changes to the composition of the community between the treated and untreated columns. Taken together, our results support the hypothesis that hypophosphite can intercept syntrophic formate exchange and that long-term treatment with hypophosphite redirects electron flow towards electron carriers other than formate.

## Results

### Start-Up performance of UASB columns

To evaluate the impact of hypophosphite on anaerobic digestion of complex organic feedstocks, we constructed six bench-scale UASB columns; three were designated for hypophosphite treatment and three served as untreated controls (**Figure S1**). We inoculated the columns with anaerobic digestion sludge from a water treatment facility and fed them whey, a common waste stream from cheese production (Pires et al., 2021). Column temperatures were maintained at 19.9°C (±0.7°C) throughout operation. To prevent souring from the difference in growth rates among microorganisms involved in fermentative methanogenesis, we gradually increased the columns feed in a stepwise manner by adjusting either the influent whey concentration or flow rate (Suryawanshi et al., 2013). Whey concentrations gradually increased in the influent from 1 g COD whey/L to 4 g COD whey/L (**Figure 1A**). Increases in whey concentration or flow rate only occurred when whey transformation stabilized. We found column whey transformation was stable when 60–70% of influent soluble chemical oxygen demand sCOD) was removed relative to the effluent sCOD (**Figure 1B**). Soluble chemical oxygen demand removal during startup and treatment aligned with previous ∼20°C UASB studies reporting steady-state sCOD removal efficiencies between 70–85% (Singh and Viraraghavan, 1998).

**Figure 1.**
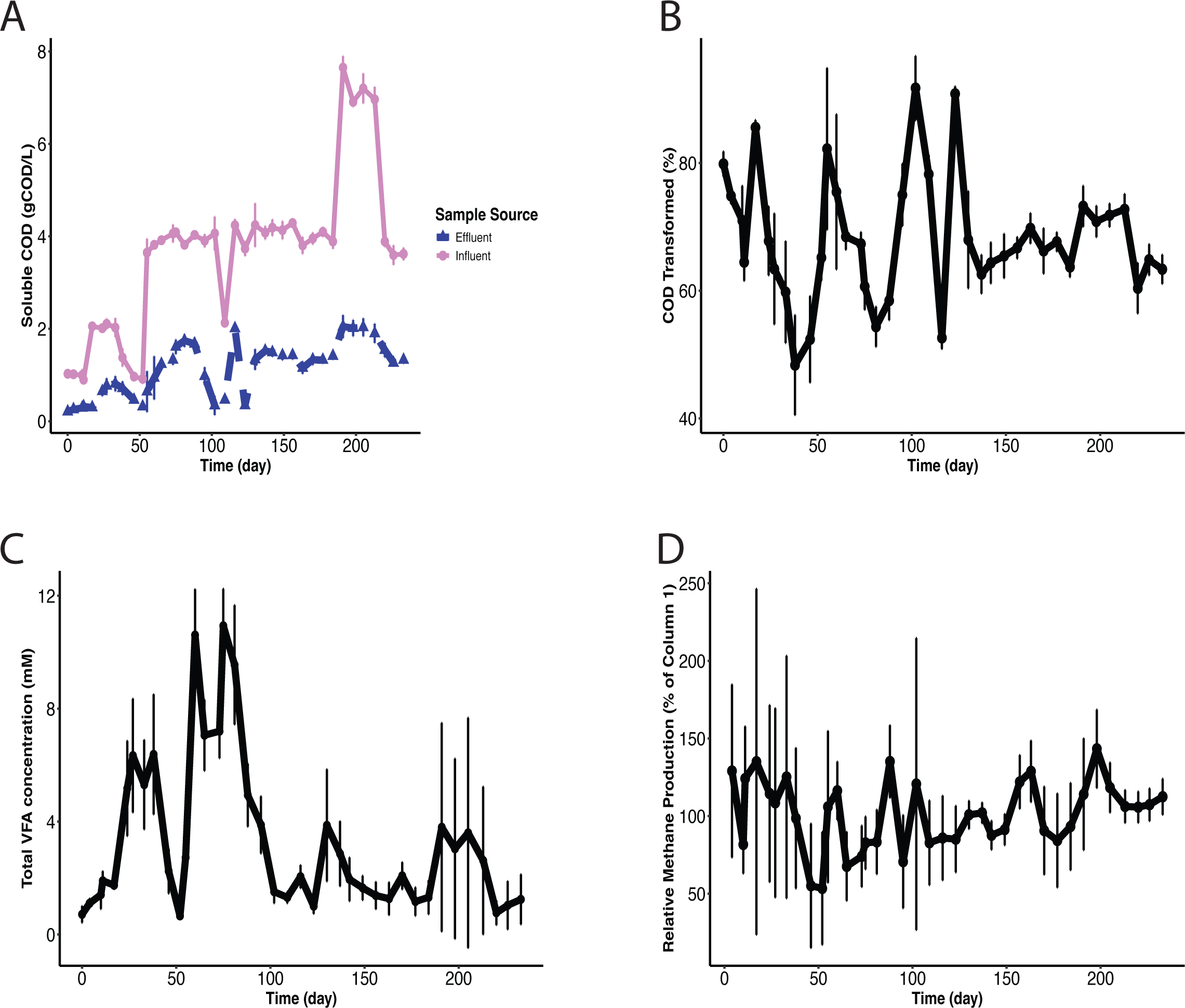
Operational parameters and performance of the UASB column during startup. **(A)** Influent (violet, solid lines and circle) and effluent (blue, dashed lines and triangles) soluble chemical oxygen demand (sCOD) over the 233-day start up. (**B)** Percent of sCOD transformed from the influent to effluent lines. (**C)** Total measured volatile fatty acid (VFA) concentrations. (**D)** Relative methane production across all columns normalized to column 1. (**A-D**) For all panels, the error bars represent the deviation from all six columns (n=6).

We monitored concentrations of lactate and seven commonly produced volatile fatty acids (VFAs) – formate, acetate, propionate, butyrate, isobutyrate, valerate, and isovalerate – to monitor primary fermentation and ensure that there was no over production of toxic VFAs. Around day 50, the pH hovered between 8 to 8.5 and the total VFA concentrations reached ∼10 mM. The accumulation consisted primarily of acetate (∼6 mM) and propionate (∼4 mM), with micromolar concentrations of butyrate, isobutyrate, valerate, and isovalerate (**Figure 1C; Figure S2**). This uncoupling between primary fermentation and syntrophic methanogenesis aligns with previous observations of reduced methanogenic activity under alkaline conditions (Qiu et al., 2023). During the weeks around day 50, the propionate: acetate ratio exceeded 1.4:1, indicating a stressed and unproductive system (D. T. Hill et al., 1987). Otherwise, the ratio remained below 1:1, reflecting a stable and productive fermentative methanogenic system (D. T. Hill et al., 1987; Hill and Holmberg, 1988; Mei et al., 2019). To help stabilize pH and reactor performance after 10 mM VFAs accumulated, we added a 40 mM HEPES buffer to adjust the pH to 7.05 at day 87, and the columns were operated in batch mode until propionate concentrations decreased to approximately 1 mM.

Along with sCOD and VFAs, we monitored methane to ensure comparable production across all six columns. During the first 120 days of operation, methane production variability between the six columns was high with some producing twice as much as others (**Figure 1D**). This high variability coincided with the alkaline pH and build-up of VFAs, both of which are known to suppress methane production (Hill et al. 1987; Qiu et al. 2023). However, after stabilizing the pH with HEPES buffer, the methane concentrations stabilized and were consistent across all six USAB columns for the remainder of the startup phase prior to treatment addition. We concluded the startup period on day 233 when no measurable differences were observed among columns in the whey transformation efficiency, VFA profiles, methane fraction, and pH, as well as acetate concentrations stabilized near 1 mM. The hypophosphite treatment period was defined as beginning on day 239 (first day), and hypophosphite dosing commenced at the end of day 244 (fifth day).

### Hypophosphite disrupts transformation of whey into methane

Based on our previous work (Hu et al., 2026), we hypothesized that the addition of 100 µM hypophosphite would be sufficient to decrease the CH4 generated from the whey. To test this hypothesis, we fed 100 µM hypophosphite through three of the six influent lines on day five of the treatment period (or day 244 of operation). The day after the hypophosphite addition to the media, the headspace CH4 concentration of the treated columns decreased. On day eight, when hypophosphite concentrations in the treated columns reached approximately 100 µM, CH4 concentrations in the treated columns were 20–30% lower than those in the untreated controls (**Figure 2A**). This observation is consistent with previous work examining the contribution of formate-utilizing methanogenesis in anaerobic digesters, which estimated from the modified Anaerobic Digest Model No 1 (ADM1) that formate conversion can account for approximately 28–34% of methane production in anaerobic digestion systems (Sun et al., 2021). At peak inhibition on day 11, CH4 concentrations in the treated columns were 20–50% lower than concentrations in the untreated columns (**Figure 2A**). Following day 11, we observed CH4 concentrations in the column headspace gradually increase in the treated columns until returning to concentrations measured in the untreated controls (**Figure 2A**), indicating an adaptation of the microbial community. Our results are consistent with our group’s previous work demonstrating amendments of sub-millimolar concentrations of hypophosphite are sufficient to reduce methane production by interfering with syntrophic formate exchange (Hu et al., 2026).

**Figure 2.**
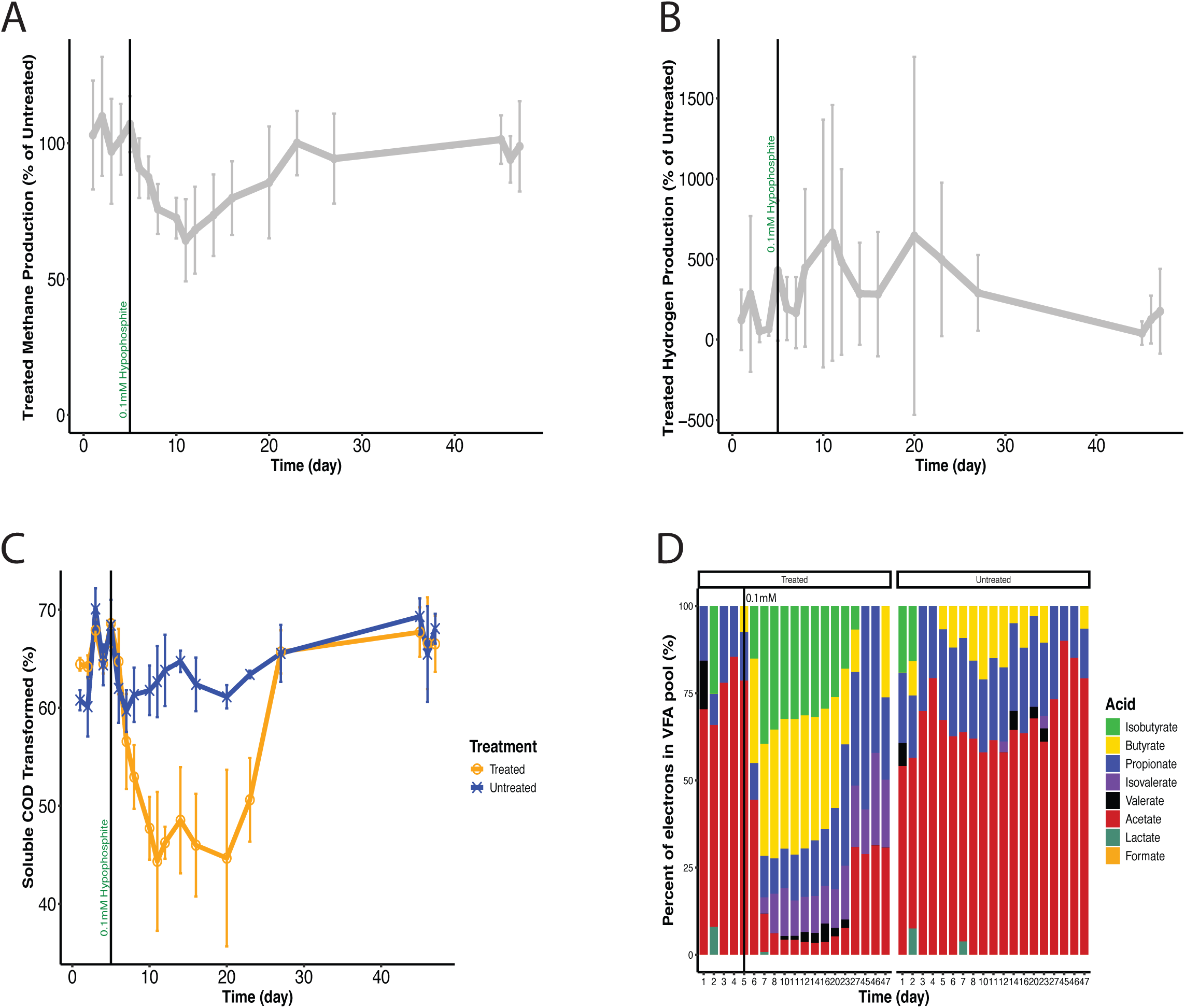
Effects of initial dosing of hypophosphite on UASB column performance. **(A)** Percentage of methane accounted for in the treated columns relative to the untreated columns (mMeffluent/mMinfluent*100). (**B)** Percentage of hydrogen gas accounted for in the treated columns relative to the untreated columns (Paeffluent /Painfluent*100). (**C)** Percent of sCOD transformed from the influent to effluent lines. The blue lines represent the columns designated as the untreated controls, and the orange lines represent columns designated for treatment with hypophosphite. (**D)** Average of relative contribution of electrons associated with individual VFAs, expressed as a percentage of the total VFA-measured electrons for the treated (n=3) and untreated (n=3) columns. (**A-C)** The error bars for treated (n = 3) and untreated (n = 3) represent standard deviation. The black line at day five represents the evening that 0.1mM hypophosphite began amendment into the treated columns.

Concomitant with the hypophosphite-dependent decrease in methane production, H2 concentrations increased in the treated columns. Of the three treated columns, two exhibited significant increases in H2 concentrations after hypophosphite addition on day 5 (p < 0.05, Welch’s two-sample *t*-test) (**Figure S3A**). On day 12, the average relative hydrogen partial pressure in all three of the treated columns reached concentrations 4.83-fold greater than the concentration in untreated columns (**Figure 2B**). The concentrations of H2 during this period surpassed the theoretical threshold for syntrophic propionate and butyrate oxidation to occur (Boone et al., 1989). Following day 12, H2 concentrations began dropping until day 45 when H2 returned to concentrations comparable to the untreated controls (**Figure 2B**). The rise increase in H2 concentrations following inhibition of syntrophic-methanogenic formate exchange by hypophosphite is consistent with H2 trends from engineered inoculum that directly inhibit methanogenesis, and indirectly syntrophy, by using methanogen-specific inhibitors (e.g., BES) (Braga et al., 2016). It also aligns with observations in *Clostridium thermocellum* cultures, where hypophosphite caused a hydrogen increase without affecting final biomass (Rydzak et al., 2014).

Prior to hypophosphite treatment, transformation of the influent whey sCOD averaged 64.6% (±3.9%) across all columns (**Figure 2C**). We maintained influent whey concentrations at 4 g COD/L (±0.22 g COD/L) throughout the time course for both treated and untreated columns. Effluent sCOD concentrations were similar among all columns, averaging 1.4 g COD/L (±0.12 g COD/L) (**Figure S3B**). By day eight of the treatment period, significantly less influent sCOD was transformed in the treated columns relative to the untreated controls (p < 0.05, Welch’s two-sample *t*-test) (**Figure 2C**). Effluent sCOD concentrations in the treated columns continued to rise until days 11–12 before levelling off at ∼20% reduction in the efficacy of whey transformation relative to the untreated controls (**Figure 2C**). Effluent sCOD in the treated columns remained approximately 33% higher than in the control columns until day 20, after which sCOD concentrations gradually declined, returning to pre-treatment levels by day 27.

The observed increase in the treated columns effluent sCOD could not be attributed to the chemical reactivity of hypophosphite in the COD assay, as the maximum g COD/L that 0.1mM hypophosphite could contribute was 49 µg COD/L. Similarly, at 71 mg Cl⁻/L, chloride concentrations are 10-fold below the chloride ion concentration that impacts COD measurements (Boyles, 1997). Therefore, the elevated sCOD indicates accumulation of reduced carbon within the treated columns, likely as unfermented whey or partially fermented intermediates. Together, these results show that sub-millimolar hypophosphite disrupted CH4 production and whey transformation for nearly three weeks before they returned to pre-treatment levels.

### Hypophosphite treatment alters VFA pools

To determine the composition of the effluent sCOD, we measured concentrations of lactate and seven individual VFAs. Prior to hypophosphite addition to the treated columns, acetate dominated the organic acid pool with concentrations ranging from 0.5–1.5 mM. Following hypophosphite addition on day five, butyrate, isobutyrate, and propionate began to accumulate in the treated columns, reaching concentrations of 2.8 mM (±0.50 mM), 2.37 mM (±0.71 mM), and 1.37 mM (±0.81 mM), respectively (**Figure S4**). Isovalerate and valerate accumulated between 300-600 µM following the increase of isobutyrate, butyrate, and propionate, while formate and lactate concentrations remained below the limit of detection. The lack of detectable formate is consistent with its role as a transient electron carrier in stable fermentative methanogenic cultures at equilibrium, as methanogens typically maintain formate at low micromolar concentrations (Schink et al., 2017).

During this period, the ratios of butyrate, isobutyrate, and propionate relative to acetate all exceeded 2:1, with isobutyrate:acetate and butyrate:acetate reaching ratios of 3.93 (±2.33) and 4.73 (±2.51) on day 12, respectively. Elevated propionate:acetate ratios are typically indicative of metabolic stress or process imbalance in anaerobic digestion systems (D. T. Hill et al., 1987; Hill and Holmberg, 1988). Notably, acetate concentrations did not increase during the treatment period, suggesting that acetoclastic methanogenesis was not directly impacted. This interpretation is consistent with previous work demonstrating that pure culture acetoclastic methanogens are not inhibited by hypophosphite until concentrations approach 100 mM (Huser et al., 1982).

As total VFA concentrations in the treated columns began to decline around day 16, propionate and isovalerate concentrations remained elevated longer than butyrate and isobutyrate. The prolonged elevation of propionate concentrations agrees with previous studies showing that propionate oxidation is more sensitive to formate and hydrogen compared to butyrate degradation (Boone et al., 1989; McInerney et al., 2008). In addition, other studies report that syntrophic isovalerate and propionate degraders often exchange electrons through formate with their methanogenic partners (Dong and Stams, 1995; Sieber et al., 2014; Chen et al., 2020). Although methane production recovered in the treated columns, the VFA concentrations remained elevated in both relative abundance and absolute concentration compared to pre-treatment conditions and the control reactors throughout the entire treatment period (**Figure 2D; Figure S4A**). This persistent elevation of VFA concentrations suggests persistent stress on the syntrophs, which serve as the primary sinks for VFA degradation in these systems (McInerney et al., 2008).

While total VFA concentrations increased in the treated columns following the addition of 100 µM hypophosphite, acetate concentrations remained stable **(Figure S4A).** Prior to hypophosphite addition, acetate accounted for more than 70% of the total VFA electron equivalents (**Figure 2D; Figure S4B**). Following treatment, the fraction of electrons associated with acetate decreased to below 8% as concentrations of propionate, butyrate, isobutyrate, valerate, and isovalerate increased. Accumulation of these VFAs resulted in the treated columns’ pH decreasing (**Figure S5A**), consistent with work demonstrating organic acids with pKas below 5 acidify their environment (Sun and O’Riordan, 2013; Cheah et al., 2019; Li et al., 2022). However, the HEPES buffer in the feed limited the magnitude of the pH change to only 0.2 pH units. As with the gas production and sCOD, by day 27 the VFA pools in the treated columns began to return to pre-treatment levels. Butyrate and isobutyrate fell from ∼75% of the total VFA electron equivalents to ∼20%, which coincided with the pH recovery to neutrality (**Figure 2D; Figure S5A**).

To evaluate whether the increase in VFAs was the source of the increased effluent sCOD, we examined the relationship between total VFA concentrations and effluent sCOD. In the treated columns, we observed a significant positive correlation between VFA concentrations and effluent sCOD (p < 0.001; ordinary least squares). We then converted VFA concentrations to g COD/L to determine their contribution to the effluent sCOD. The difference between the non-VFA portions of the effluent sCOD between the treated and untreated columns was negligible, consistent with the continuation of primary fermentation while downstream syntrophic or methanogenic conversion was temporarily impaired (**Figure S5B**). After recovery in the treated columns, sCOD gradually decreased as VFA concentrations decreased. Overall, our results are consistent with previous studies demonstrating that hypophosphite can alter product distribution (Rydzak et al., 2014; Hu et al., 2026).

### Hypophosphite-treated columns are insensitive to higher concentrations of hypophosphite after recovery

The changes in VFA concentrations demonstrate the ability of hypophosphite to significantly disrupt formatotrophic methane production in a continuous flow system prior to adaptation and rebound. To determine if higher doses of hypophosphite would lead to further disruption, we increased the influent hypophosphite concentration in the treated columns from 0.1 mM to 1 mM on day 47 and again to 10 mM on day 53. Notably, even with hypophosphite concentrations increased to 1 mM or 10 mM, we did not observe a significant change in the amount sCOD transformed in the column, nor the gas produced (p > 0.05, Welch’s two-sample *t*-test) (Figure 3A-C. However, VFA pools remained shifted with persistently higher concentrations of VFAs in the treated columns (**Figure 3D**).

**Figure 3.**
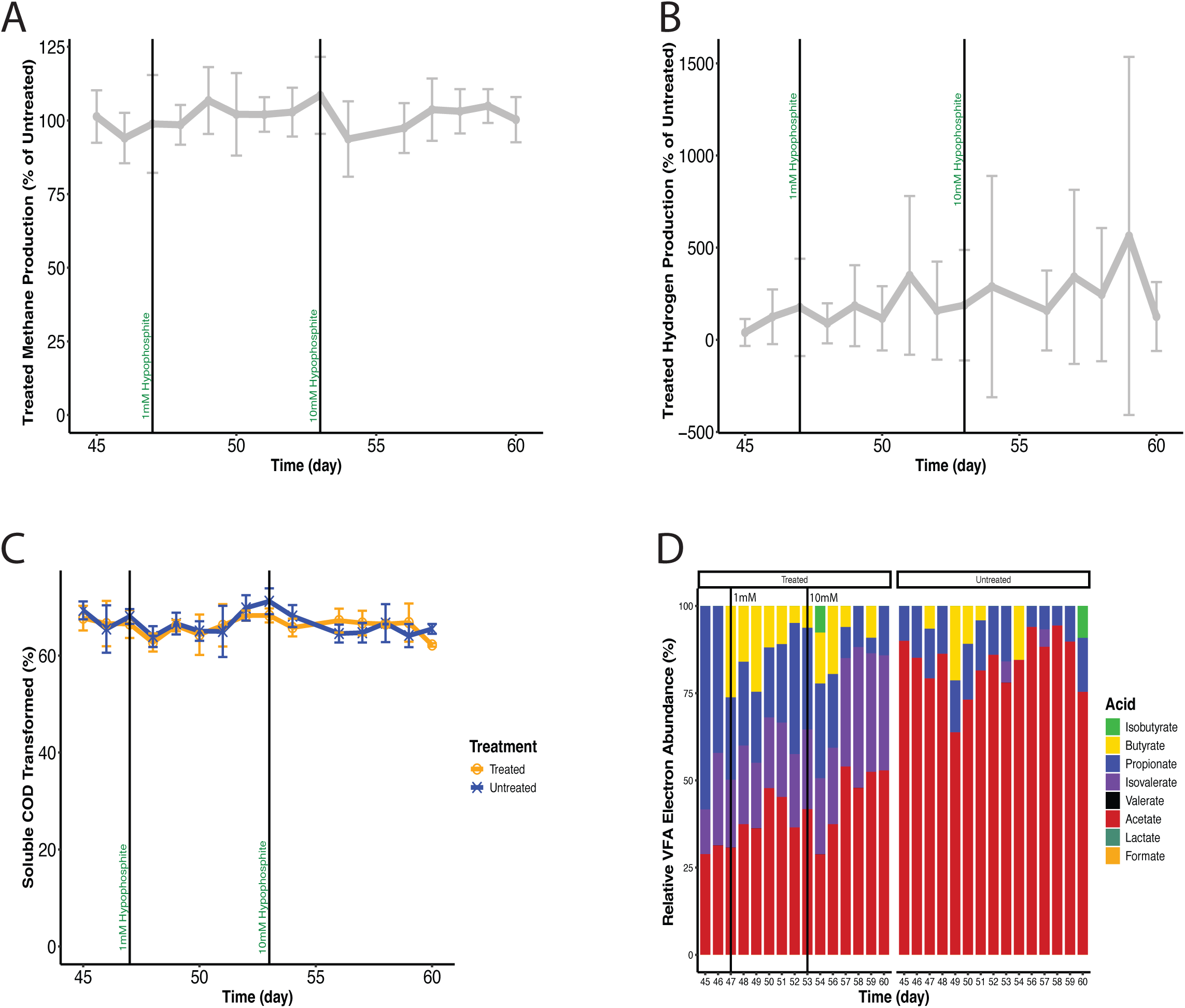
Addition of 1 and 10mM hypophosphite does not alter column parameters. **(A)** Percentage of methane accounted for in the treated columns relative to the untreated columns (mMeffluent/mMinfluent*100). (**B)** Percentage of hydrogen gas accounted for in the treated columns relative to the untreated columns (Paeffluent /Painfluent*100). (**C)** Percent of sCOD transformed from the influent to effluent lines. The blue lines represent the columns designated as the untreated controls, and the orange lines represent columns designated for treatment with hypophosphite. (**D)** Average of relative contribution of electrons associated with individual VFAs, expressed as a percentage of the total VFA-measured electrons for the treated (n=3) and untreated (n=3) columns. **(A-C)** The error bars for treated (n = 3) and untreated (n = 3) represent standard deviation. The black line at day 47 represents the evening that 1 mM hypophosphite began amendment into the treated columns. The black line at day 53 represents the evening that 10 mM hypophosphite began amendment into the treated columns.

### Microbiome compositions of treated columns are not significantly different from untreated columns

To determine if the changes in VFA composition and gas production could be attributed to a shift in the microbiome, we used 16S rDNA amplicon sequencing to measure the microbiome composition of the UASB columns over the course of the column operation. The sludge communities were primarily composed of nine phyla: Firmicutes, Bacteroidota, Halobacterota, Spirochaetota, Synergistota, Desulfobacterota, Proteobacteria, Euryarchaeota, and Actinobacteriota. Firmicutes, Bacteroidota, and Halobacterota emerged as the dominant phyla by relative abundance (**Figure 4A; Figure S7A**). Abundant genera within the phylum Bacteroidota included *Bacteroides*, *Dsygonomonas*, *Macellibacteroides*, and *Williamwhitmania*. Of these, *Bacteroides* was the most prevalent and isolates of this genus are known to ferment sugars and some amino acids to produce H2, formate, and organic acids such as pyruvate, propionate, acetate, lactate, succinate, butyrate, isobutyrate, and isovalerate (Frantz and McCallum, 1979; Rios-Covian et al., 2017; Shin et al., 2024). Similarly, the core Firmicutes genera – *Clostridium sensu stricto 1*, *Eubacterium*, *Trichococcus,* and *Acidaminococcus* – can utilize a range of carbohydrates and some amino acids to produce acetate, formate, H2, propionate, butyrate, isobutyrate, lactate, and alcohols (Liu, 2002; Duncan et al., 2004; Jumas-Bilak et al., 2007; Engels et al., 2016; Li et al., 2023). Within Halobacterota, *Methanosaeta* and *Methanocorpusculum* represented much of the phylum. Previous work demonstrated *Methanosaeta*’s ability to generate methane from acetate or direct interspecies electron transfer (DIET), while *Methanocorpusculum* converts H2 or formate to CH4 (Zellner et al., 1987; Ma, 2006; Rotaru et al., 2014; Volmer et al., 2023).

**Figure 4.**
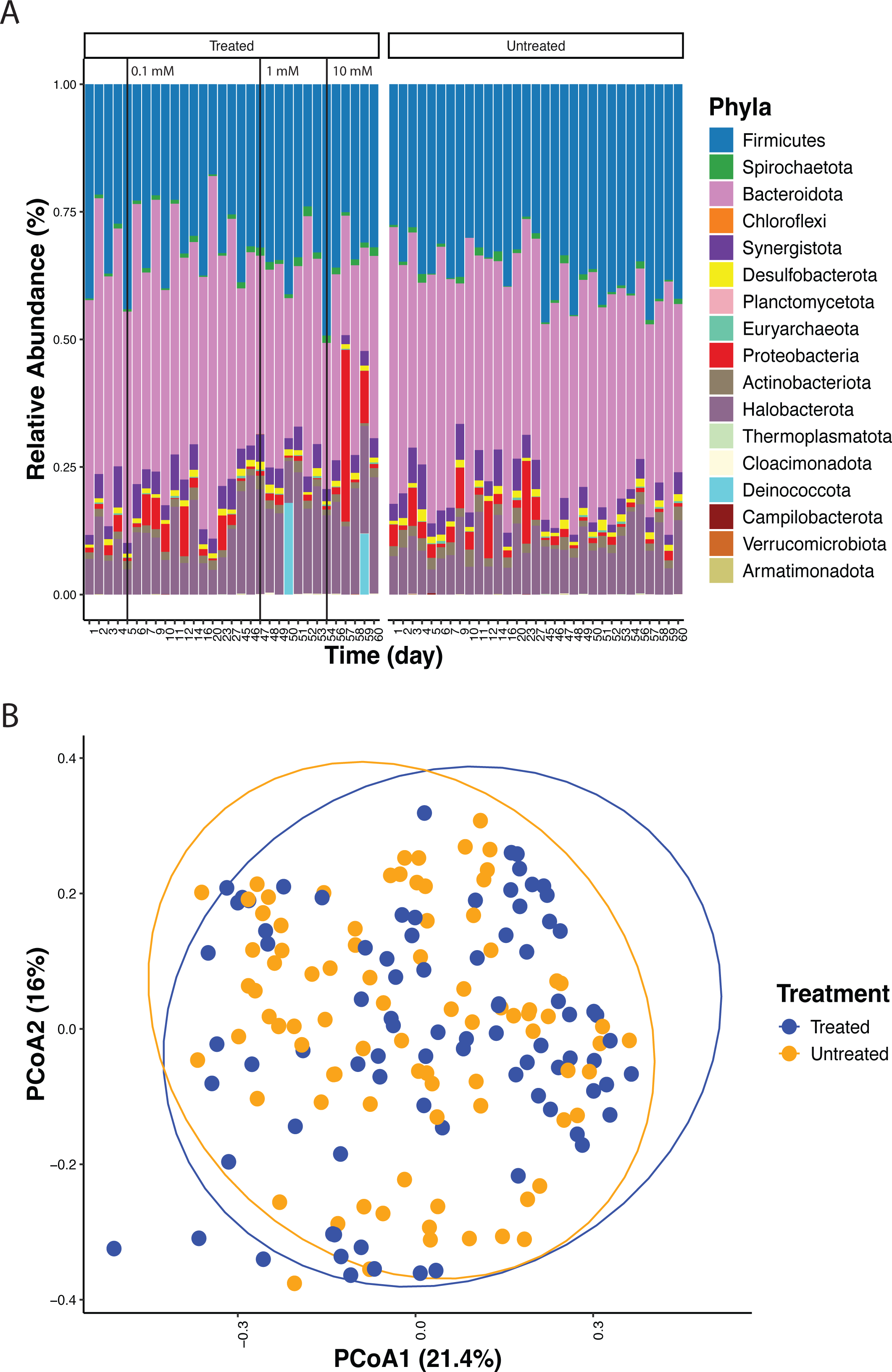
UASB microbiome composition over time in treated and untreated columns. (**A)** Average of relative abundance of prokaryotic phyla over the course of the UASB operation. The black line at day five represents the evening that 0.1 mM hypophosphite began amendment into the treated columns. The black line at day 47 represents the evening that 1 mM hypophosphite began amendment into the treated columns. The black line at day 53 represents the evening that 10 mM hypophosphite began amendment into the treated columns. (**B)** Principal component analysis comparing treated and untreated columns.

Diversity and evenness calculations indicated that the addition of hypophosphite did little to alter the microbiome composition over time (**Table S1,S2**). Specifically, Shannon’s Diversity, Simpson’s Diversity, and Richness scores were consistent over time across all columns (Table S1, S2). A few time points differed significantly throughout the time course; though, none coincided with hypophosphite dosing and were transient (Kruskal-Wallis and Dunn’s post-hoc, p < 0.05). To determine whether hypophosphite treatment impacted the microbiome, we compared treatment groups using Bray-Curtis dissimilarity. Microbiome composition did not differ significantly between treatments (PERMANOVA, F =3.08, R^2= 0.016, p =0.20) (**Figure 4B**), and dispersion was homogenous between groups (PERMDISP, F = 0.32, p = 0.58), indicating the lack of significance was not confounded by unequal within-group variation. To determine if a specific time point might be significant, Bray-Curtis dissimilarity was calculated again but between the treatment groups at each time point; however, no single time point differed significantly between treatment groups (PERMANOVA, p > 0.05) (**Table S1**). Thus, changes in microbiome composition do not explain the changes in VFA composition and gas production at 0.1 mM hypophosphite, or the lack of response at 1 and 10 mM hypophosphite (**Figure 2-4**).

### Formatotrophic methanogenic activity is reduced in treated column biomass

To assess the capacity of the active sludge biomass in the UASB columns to utilize formate or H2 as a methanogenic substrate, biomass suspensions from the columns were pooled within each treatment, washed to remove residual hypophosphite and metabolites and incubated with either formate or hydrogen as an electron donor. We found that while the capacity for hydrogenotrophic methanogenesis persisted in the treated columns’ sludge, formatotrophic methanogenesis was significantly reduced (p < 0.05, t-test) (**Figure 5A**). Formatotrophic methanogenesis rates were ∼36 times higher in the untreated sludge compared to the treated sludge (16.11 ± 2.97 mM CH4/g dry weight/hr versus 0.45 ± 0.05 mM CH4/g dry weight/hr). Hydrogenotrophic methanogenesis rates remained comparable across treatments (p > 0.05, t-test). However, for the treated sludge, CH4 production rates were nearly 10-times higher in the H2 condition than the formate-amended condition (4.41 ± 0.56 mM CH4/g dry weight/hr versus 0.45 ± 0.05 mM CH4/g dry weight/hr; p < 0.05, t-test). Consistent with this reduced formatotrophic capacity, formate-amended treated samples exhibited the longest delay before CH4 production significantly surpassed the baseline level (p < 0.05, t-test) (**Figure 5B**). This metabolic shift away from formate utilization by the microbiome aligns with both prior studies and our own work, suggesting that hypophosphite suppressed formate production and consumption (Takamiya, 1953b; Pine and Vishniac, 1957; Pine, 1958; Gründig and Babel, 1987; Rydzak et al., 2014; Lee et al., 2025; Hu et al., 2026).

**Figure 5.**
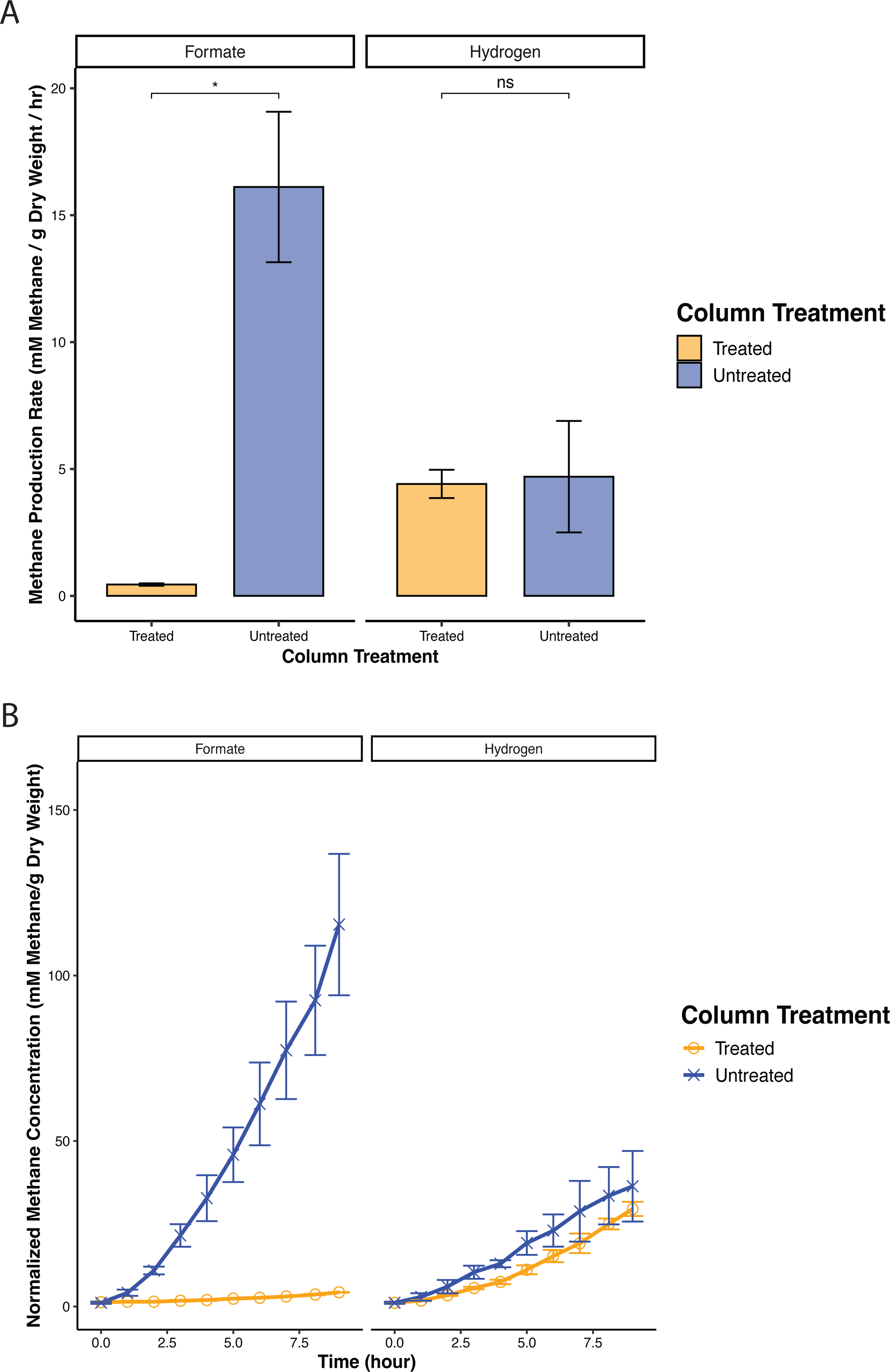
Hydrogenotrophic versus formatotrophic methanogenic activity in treated and untreated columns. **(A)** Methane production rates (mM/g dry weight/hr) from the treated and untreated washed sludge activity assays amended with either hydrogen or formate. (**B)** Normalized methane concentration (mM/ g dry weight) over time. “*” = p < 0.05. “**” = p <0.01.

## Discussion

Our objective was to determine the influence of the formate analog, hypophosphite, on whey-degrading fermentative methanogenic continuous-flow UASB columns. We demonstrate that sub-millimolar concentrations are sufficient to significantly reduce methane concentrations by ∼30%, while redirecting carbon and electron flow toward the accumulation of VFAs and hydrogen (**Figure 2**). Following the initial hypophosphite amendment, concentrations of propionate, butyrate, isobutyrate, valerate, and isovalerate increased rapidly, producing VFA-to-acetate ratios not observed in the untreated columns, or even during the startup phase of the columns (**Figure S2A; Figure S4A**). This observation is consistent with the hypothesis that hypophosphite selectively disrupts formate-mediated interspecies electron transfer leading to accumulation (Hu et al., 2026). Furthermore, prolonged exposure to hypophosphite concentrations ranging from 0.1 to 10 mM resulted in a reduction in formate utilization by the methanogenic sludge, with formatotrophic CH4 production rates decreasing by approximately 36-fold on average relative to un-treated sludge (**Figure 5A**). Our results support a model whereby hypophosphite redirects carbon and electron flow in fermentative methanogenic systems.

Beyond supporting the selective interference of hypophosphite with syntrophic formate exchange at low concentrations, in this study we examined the impact of continuous hypophosphite treatment on a methanogenic microbiome. The recovery of methane production following initial inhibition, the absence of shifts in microbiome composition, and the outcomes of the terminal activity assays implicate metabolic plasticity of syntrophic electron flow, with the microbiome shifting toward greater reliance on hydrogen or acetate over formate as an electron carrier **(Figure 6**). This stable influence of a metabolic inhibitor on complex carbon and electron flow opens up new possibilities. For example, combining hypophosphite with inhibitors of acetoclastic or hydrogenotrophic methanogenesis could represent new methane control strategies.

**Figure 6.**
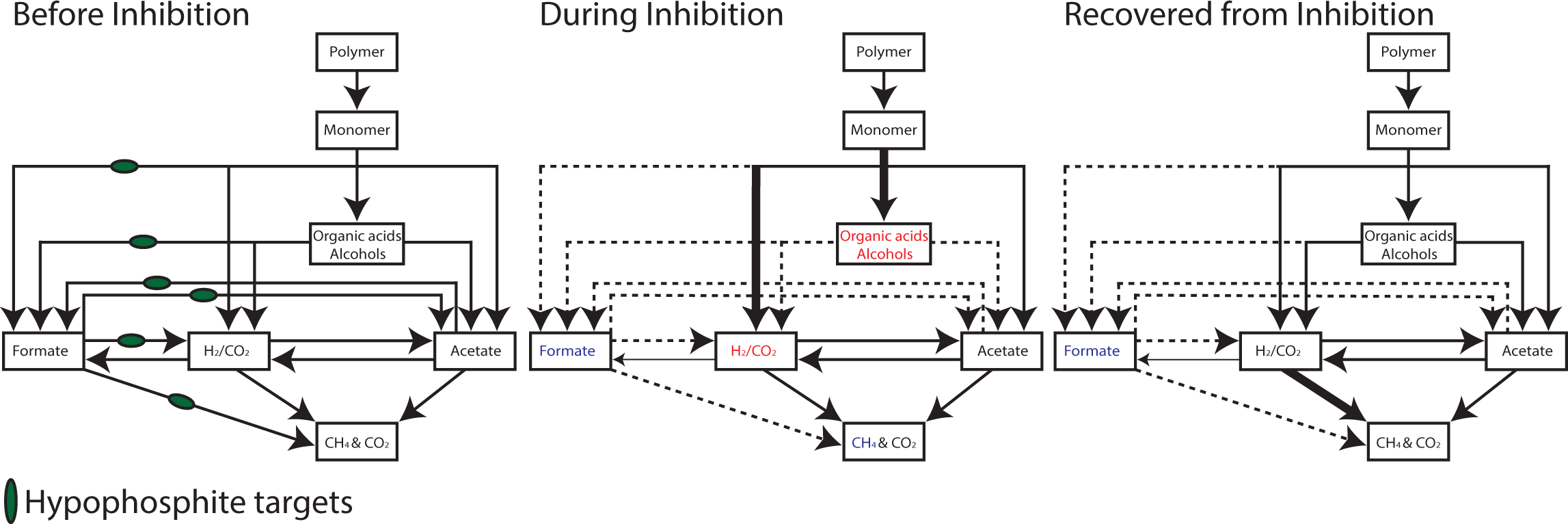
Proposed model for electron and carbon flow under hypophosphite inhibition and recovery. Schematic of electron and carbon flow in the treated columns prior to hypophosphite addition, during max inhibition, and following recovery. Solid lines represent active carbon and electron flow, whereas dashed lines represent a reduction in flow. Line thickness reflects relative flux magnitude. Color denotes metabolic concentration trends: blue, decreased; black, unchanged; red, increased.

Apart from targeting syntrophic formate exchange, higher concentrations of hypophosphite (1-10 mM) may impact production of formate by fermentative bacteria. Studies on primary fermenters have shown that amendments of hypophosphite at 1 mM are sufficient to stop formate production, and in some cases increase H2 production, without influencing growth (Rydzak et al., 2014; Lee et al., 2025). Gründig and Babel (1987) showed that a 1-minute incubation period of 12 mM hypophosphite reduced *Acetobacter methanolicus* MB58’s ability to oxidize formate during methanol and formaldehyde oxidation. Thus, in our study, hypophosphite may be redirecting the electron flux from formate to H2 by primary fermenters, alongside inhibiting the exchange of formate between syntrophs and methanogens.

Future work will focus on further characterizing the mechanisms responsible for the redirection of carbon and electron flow in our laboratory scale anaerobic digestors by hypophosphite. 16S rDNA amplicon sequencing alone cannot resolve the physiological function of each taxon or identify adaptive mutations. Thus, through shotgun metagenomics, we will aim to clarify whether any genetic mutations can explain the mechanism by which hypophosphite shifts carbon and electron flow away from formate and towards other interspecies electron carriers. Other than a study that reported a mutation in the *E. coli* formate transporter FocA during adaptive lab evolution to hypophosphite, little is known about how microorganisms may overcome hypophosphite stress or inhibition (Suppmann and Sawers, 1994). Finally, combining hypophosphite treatment with approaches for modulating the activity of fermentative methanogenic microbiomes could lead to new strategies for diverting carbon and electrons away from methanogenesis in anaerobic digestors, cattle rumens, rice fields and other methanogenic industrial ecosystems.

## Materials and Methods

### Column setup and operation

The up-flow anaerobic sludge blanket (UASB) continuous-flow experiment was run using six laboratory-scale columns (200mL). Each column was filled with 120mL of anoxic <u>B</u>icarbonate <u>W</u>hey <u>H</u>EPES (BWH) media and 8 mL of anaerobic digested sludge from Silicon Valley Clean Water Wastewater Treatment Plant, Digester 3. Anoxic media was made by adding all the components to a 1.6L bottle and then using 1M NaOH or HCl to adjust the pH to 7.05. The media was then boiled and put under a stream of N2 to cool before capping with a butyl rubber stopper. The media was sterilized in the autoclave (121°C for 45 minutes). For 4gCOD Whey/L BWH the recipe was (per L of deionized water): 3g sodium bicarbonate, 9.53g HEPES buffer, 5g bovine milk whey (Sigma Aldrich), and 10mL each of vitamin and mineral solutions described in (Figueroa et al., 2018)). Na2S*9H2O (1mM) was added after the media was autoclaved from a sterile stock solution. A line was connected from the top of the column to an inverted 250mL graduated cylinder filled with 100% saline-acidic (pH<4) water to capture gas (Walker et al., 2009).

After inoculation, the columns incubated for one month, during which production of methane and VFAs were monitored to assess the onset of microbial activity prior to starting continuous feeding. To stop the washout of granules and sludge, 3mL syringes (BD) were cut into 1mL length rings and added so the top 1-3 inches of the liquid layer was covered by them. All columns were started at an organic loading rate (OLR) of 0.082 gCOD Whey/L/day and ramped up over 8 months to 1.31 gCOD Whey/L/day. The cultures were kept at room temperature (20°C) in the dark. Three of the columns were designated for treatment, and the other three were kept as untreated controls. Treatment began with a gradual increase of hypophosphite into the treated columns, taking ∼3 days to reach 100µM based on the hydraulic retention time (HRT), or the time required to replace the reactor volume. The treated columns continued receiving hypophosphite amendments of 100 µM. On day 47, the hypophosphite in the treated columns feed was increased to 1mM, and by day 50, would have reached its max concentration. One full volume change of the columns under 1mM was allowed before increasing the treated columns feed to 10mM hypophosphite on day 53.

### DNA extraction and 16S rRNA gene analysis

For 16S rDNA amplicon sequencing, 2mLs of sample were taken from the sludge bed of each column at each time point and centrifuged (14,000 rcf, 4 minutes) to collect biomass, then stored at -20°C. The gDNA was extracted using the DNAeasy PowerSoil Pro Kit (Qiagen) as described in the manufacturer’s instructions. Quality and concentration of the extracted DNA were determined by Nanodrop Lite Spectrophotometer (Thermo Fischer Scientific, MA, USA). gDNA went through a PCR reaction to amplify the V4/V5 regions of the 16S rDNA gene using the 515(-Y)F and 926R primers, but with in-line dual Illumina indexes (Parada et al., 2016; Price et al., 2018). The 16S amplicons were sequenced using an Illumina MiSeq (Illumina, San Diego, CA, USA) with 2×300bp Illumina v3 reagents. Raw reads were processed as described in Hu et al. (2025). A custom taxonomic reference database was constructed for 16S rDNA gene classification using QIIME2 (version 2024.2) (Bokulich et al., 2018; Robeson et al., 2021). Analyses were performed within a dedicated QIIME2 amplicon conda environment generated from the official QIIME2 distribution (https://library.qiime2.org). Reference sequences and taxonomy were derived from the SILVA SSU Ref NR 99 database (release 138) (Chuvochina et al., 2025). To ensure compatibility with the amplified region, reference sequences were trimmed in silico to the V4/V5 region using the primers 515F (GTGYCAGCMGCCGCGGTAA) and 926R (CCGYCAATTYMTTTRAGTTT). Primer trimming was performed using the feature-classifier extract-reads function in QIIME2 with forward read orientation and parallel processing enabled. The resulting region-specific reference sequences were used to train a naïve Bayes classifier following the workflow described by the QIIME2 classifiers repository with modifications to accommodate SILVA release 138 and the V4/V5 primer set. Taxonomic assignment of amplicon sequence variants (ASVs) was performed using the trained classifier method in QIIME2, with a confidence threshold of 0.7. Taxonomic classifications were summarized and visualized using QIIME2 metadata tabulation tools.

### Analytical Techniques

Formate, acetate, lactate, propionate, butyrate, isobutyrate, valerate, and isovalerate were quantified by high-performance liquid chromatography (HPLC) on a Shimadzu LC-20 using a Biorad Aminex® HPC-87H, 300mm x 7.8mm column with a 5mM sulfuric acid mobile phase at a flow rate of 0.6mL/min. The temperature was held at 50°C. We quantified organic acids using a UV detector at 216 nm. The limit of detection was 125 µM for all acids. pH was quantified using a TS-ISE500 Lab pH/ISE meter and 2mLs of filtered (0.22µm) sample. To determine the soluble COD (gCOD/L) of the influent and effluent, 3mLs of sample were taken from the influent and the effluent ports, respectively. Samples were filtered through a 0.22µm filter and diluted by a factor of 10 in deionized water. Samples were handled as described in the kit’s manual, Chemical Oxygen Demand TNTplus Vial Test, HR (20-1,500 mg/L COD) (Hach). The chemical oxygen demand of hypophosphite was calculated to be 0.49 g CO per mole. The optical density of the columns was measured on a ThermoScientific Genesys 20 at a wavelength of 600nm. Methane was measured using an Agilent Technologies 7890a gas chromatograph flame ionization detector (GC-FID). The GC oven temperature and column (Supelco 2380, 30m x 250µm x 0.2µM, flow = 1mL/min) were heated to 50°C and held constant for analysis. 0.5mL was pulled from the headspace, then pushed to 0.1mL and injected into the GC-FID. Hydrogen in the headspace was measured using a Peak Performer 1 (PP1) hydrogen analyzer (Peak Laboratories, CA).

### Diversity Indexes and Statistical analysis

To assess microbial alpha diversity within each sample, we rarefied all samples to 2,000 sequences. Richness, Shannon diversity, and Simpson diversity indices were then calculated for each column at each time point. Normality of each alpha-diversity metric was evaluated using the Shapiro–Wilk test. Differences in alpha-diversity metrics between treated and untreated columns at each time point were evaluated using the nonparametric Kruskal–Wallis test, followed by Dunn’s post hoc test where appropriate. Statistical significance was defined as *p* < 0.05 and is indicated by an asterisk (*). To assess changes in community composition associated with hypophosphite treatment, beta diversity was calculated using Bray–Curtis dissimilarity. Comparisons were performed both across the full experimental time course and between treated and untreated columns at individual time points. Beta-diversity calculations and visualizations were based on Zotus derived from 16S rDNA gene amplicon sequencing. Prior to beta-diversity significance testing, homogeneity of dispersion between treatment groups was assessed using PERMDISP (*betadisper*). All statistical analyses and data visualization were conducted in R (version 4.5.0). All diversity metrics calculated used the *vegan* (version 2.7-1) package in R (Oksanen et al., 2025). Overall differences in community composition between treated and untreated columns were evaluated using permutational multivariate analysis of variance (PERMANOVA) in R. The six columns were treated as the independent experimental units, with repeated samples from the column over time retained as observations rather than independent treatment replicates. Treatment significance was assessed using an exact permutation test that reassigned treatment labels across all possible column-level groupings, keeping all the time points from a given column together. Pairwise comparisons at individual time points were conducted using PairwiseAdonis (Martinez Arbizu, 2020). To evaluate linear relationships between measured analytes, ordinary least squares (OLS) regressions were performed with significance defined as *p* < 0.05. Regression analyses and visualizations were generated using *ggplot2* (version 3.5.2.). Differences in individual analyte concentrations (e.g., g COD L⁻¹) between treatment conditions at specific time points were assessed using Welch’s two-sample *t*-test, with significance determined at *p* < 0.05.

### Cell Suspension preparation

After time point 60 of the hypophosphite treatment period, we collected and concentrated biomass from both untreated and treated columns from 360 mL of sludge under a continuous stream of nitrogen to maintain anaerobic conditions, separately. The resulting pellets were washed three times under nitrogen with a modified hypophosphite-free buffered minimal salts medium originally developed for the isolation of *Phosphitispora fastidiosa* DYL19 (Harris and Kline, 1956; Mao et al., 2021). The modification consisted of substituting the vitamin and mineral solutions with those described by (Figueroa et al., 2018). Following washing, we resuspended the biomass in 25 mL of the same anoxic buffered medium to generate a homogeneous cell suspension. Aliquots (1 mL) of the sludge suspension were dispensed into the same anoxic buffered media amended with either formate (25mM) or hydrogen (10mLs) for both treated and untreated samples. Methane production was measured by GC-FID as described above. Incubations were maintained horizontally at 20°C in the dark. At the end of the incubation, samples were centrifuged and stored at −20°C prior to dry weight determination. Dry biomass weight was determined by drying samples at 90°C overnight to remove residual water. Methane production was monitored over time, and rates were calculated from the linear portion of methane accumulation curves and normalized to biomass dry weight.

## Acknowledgments

We would like to acknowledge Nivriti Krishnamurthy for experimental help used during column operations. We would also like to thank Chandler Sutherland and Zachary Hallberg for feedback on the manuscript. Additionally, we would like to thank Silicon Valley Clean Water for providing the anaerobically digested sludge used as inoculum for this paper. This study was financially supported by the Energy & Biosciences Institute (EBI) through the EBI-Shell program and some co-authors are members of ENIGMA (Ecosystems and Networks Integrated with Genes and Molecular Assemblies; (http://enigma.lbl.gov), a Science Focus Area Program at Lawrence Berkeley National Laboratory, US Department of Energy, Office of Science, Biological and Environmental Research Program under contract number DE-AC02-05CH11231 to Lawrence Berkeley National Laboratory.

## Author Contributions

Conceptualization: M.E.W., R.H., H.K.C., Y.L., and J.D.C.; Investigation: M.E.W. and A.W.; Formal Analysis: M.E.W.; Visualization: M.E.W.; Funding Acquisition: J.D.C. and H.K.C; Writing – original draft: M.E.W.; Writing – review & editing: M.E.W., R.H., H.K.C., A.W., Y.L., and J.D.C.

## Data Availability

All data needed to evaluate the conclusions in the paper are present in the paper and/or the Supplementary Information. The nucleotide sequences for 16S rDNA gene amplicon sequencing and metagenomic sequencing were deposited in the SRA database under accession numbers PRJNA1434523.

## Code availability

All software packages utilized in this study are publicly accessible, and no original code is reported in this study

