## Supplementary Materials for "Redirecting carbon and electron flow in methanogenic laboratory scale anaerobic digestors with hypophosphite"

### Inhibition of methanogenic fermentation in a flow-through column by hypophosphite treatment

#### Supplementary Figure Legends

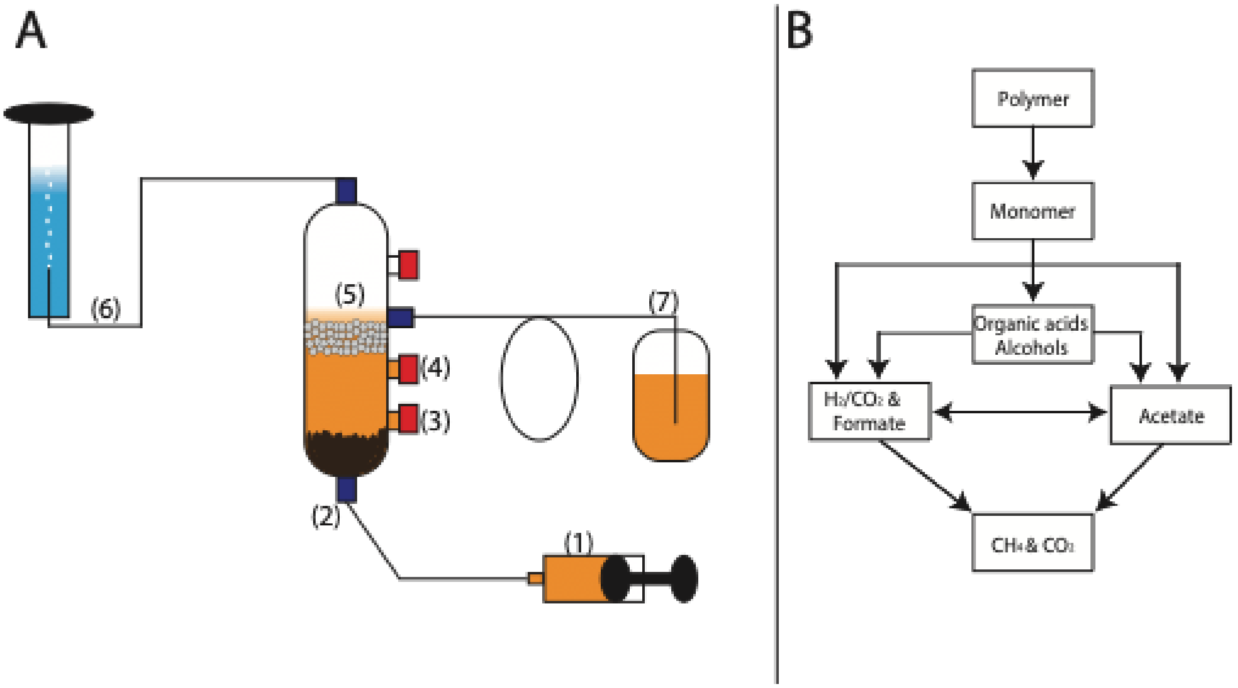


**Figure S1. Schematic of up-flow anaerobic sludge blanket (UASB) column and methanogenic complex carbon degradation. A,** (1) Syringe pump pushing feed into influent line. (2) Influent port for influent soluble chemical oxygen demand (sCOD) and 16S rRNA sequencing sampling. (3) Port for optical density ($OD_{600}$) and organic acids sampling. (4) Port for effluent sCOD and pH sampling. (5) Cut syringes to block granules and sludge from flowing out. (6) Gas-collection line and connected graduated cylinder/tub filled with 100% saline acidic (pH<4) water for gas displacement. (7) Waste collection. **B,** Model for carbon and electron flow in a methanogenic complex carbon degrading community.

A

B

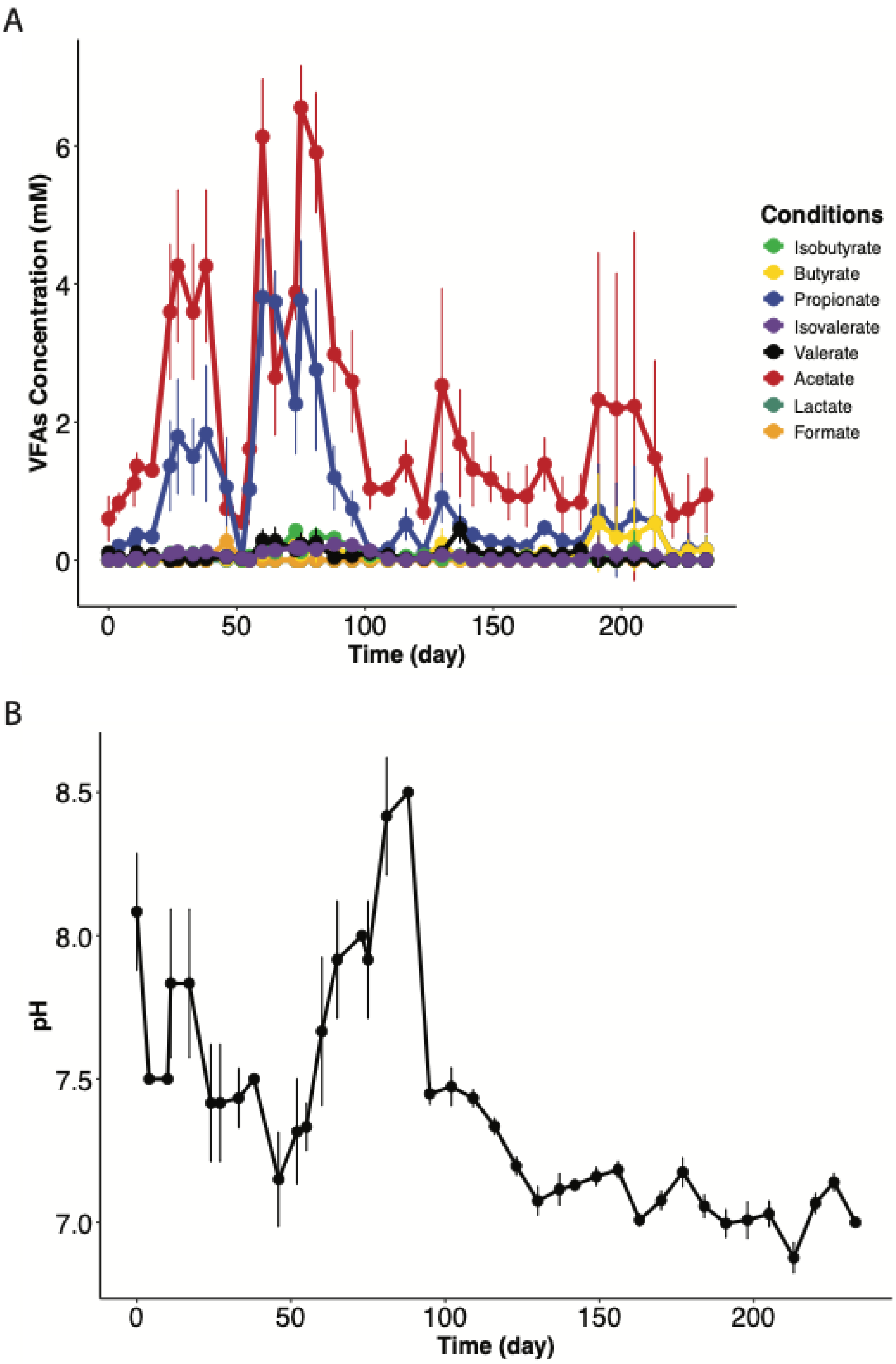

VFAs Concentration (mM)
Time (day)
Conditions
Isobutyrate
Butyrate
Propionate
Isovalerate
Valerate
Acetate
Lactate
Formate
pH
Time (day)

**Figure S2: Measured volatile fatty acid (VFA) and pH changes over column start up. A.** Concentrations of formate (orange), acetate (red), lactate (forest green), propionate (blue), isobutyrate (green), butyrate (yellow), valerate (black), and isovalerate (purple) over time during the startup of the treated and untreated designated columns. **B.** pH of treated and untreated designated columns. Error bars represent standard deviation (n=6).

A

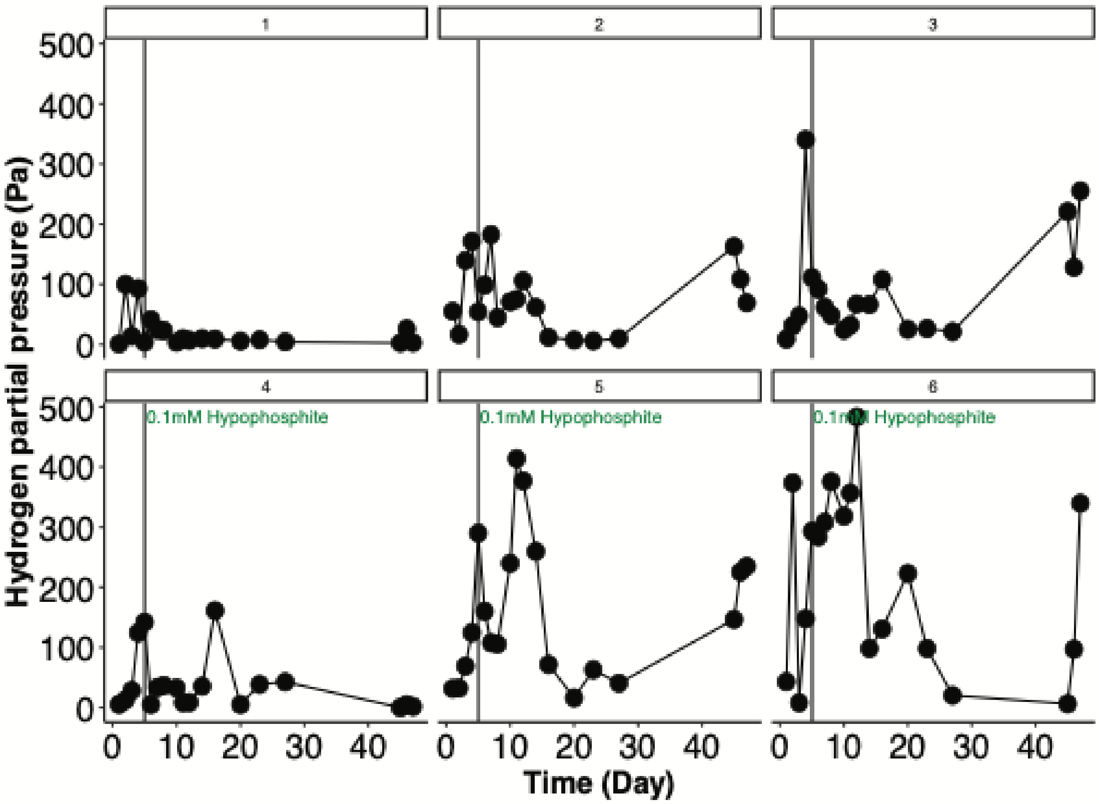
1
2
3
4
5
6
0.1mM Hypophosphite
0.1mM Hypophosphite
0.1mM Hypophosphite
Hydrogen partial pressure (Pa)
Time (Day)

B

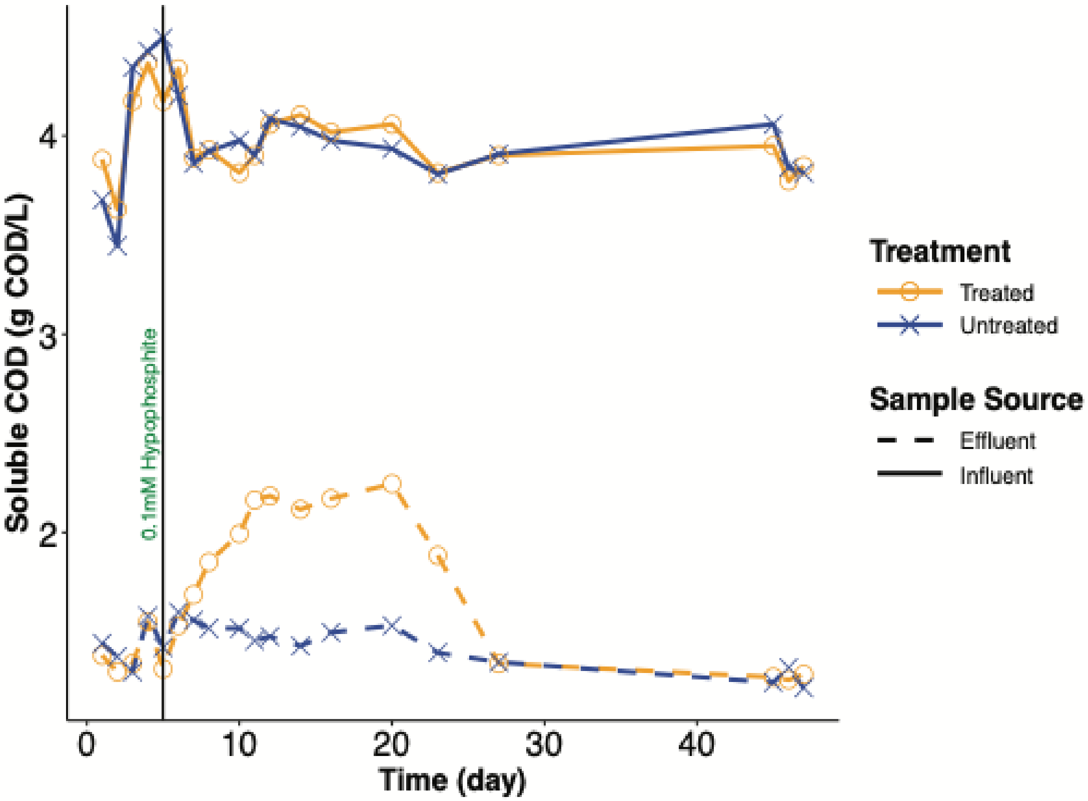
0.1mM Hypophosphite
Soluble COD (g COD/L)
Time (day)
Treatment
Treated
Untreated
Sample Source
Effluent
Influent

**Figure S3: Hydrogen ($H_2$) and soluble COD (sCOD) fluctuations during initial treatment.** **A.** Concentration of $H_2$ (Pa) in individual columns (1-3 untreated, 4-6 treated). Error bars represent standard deviation. **B.** Average concentration of sCOD (gCOD/L) found at the influent and effluent port of treated (orange) and untreated (blue) columns. Error bars represent standard deviation (n=3 per condition). The black line at day five represents the evening that 0.1mM hypophosphite began amendment into the treated columns.

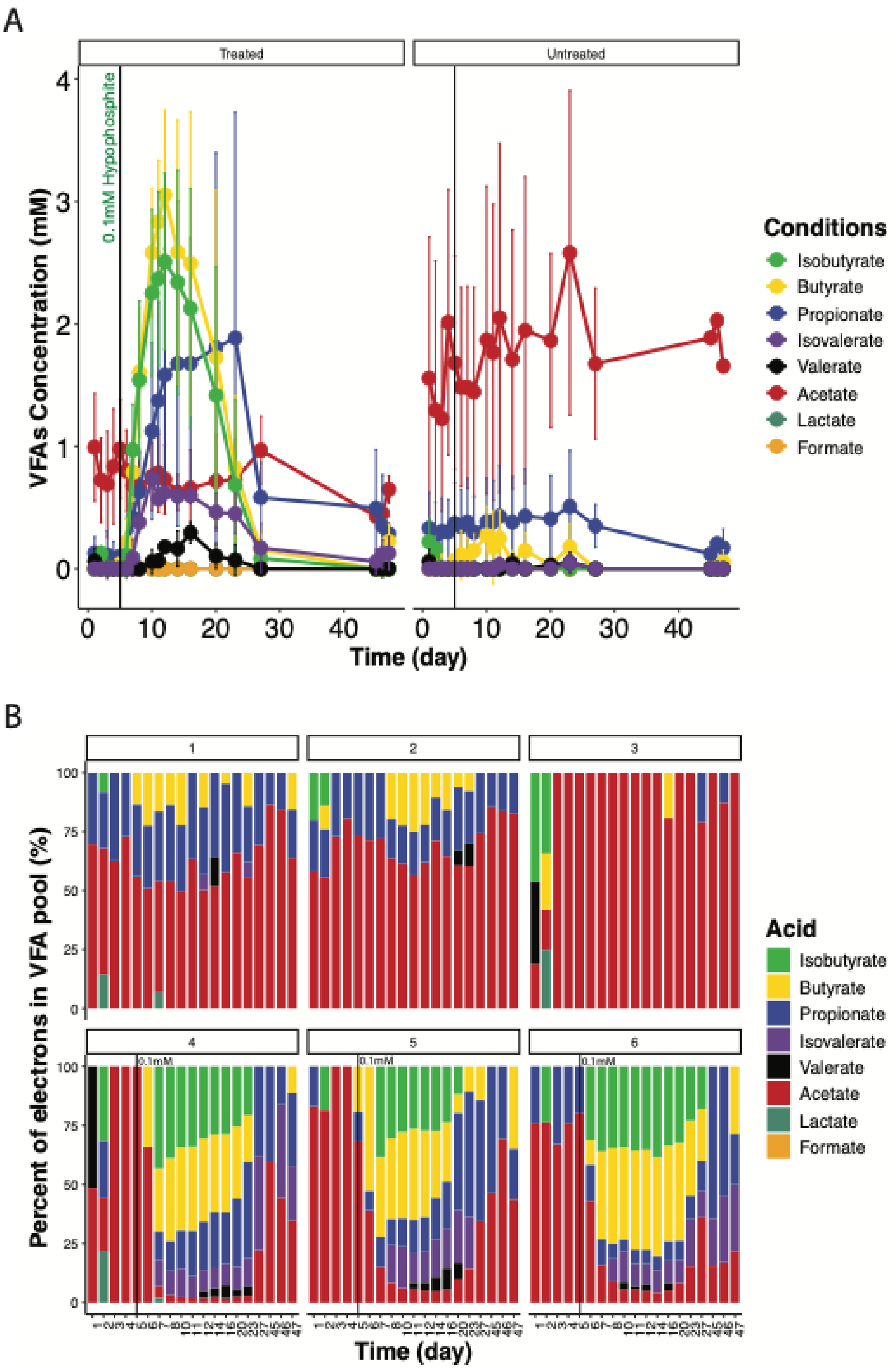
A
Treated
Untreated
0.1mM Hypophosphite
VFAs Concentration (mM)
Time (day)
Conditions
Isobutyrate
Butyrate
Propionate
Isovalerate
Valerate
Acetate
Lactate
Formate
B
1
2
3
4
5
6
0.1mM
Percent of electrons in VFA pool (%)
Time (day)
Acid
Isobutyrate
Butyrate
Propionate
Isovalerate
Valerate
Acetate
Lactate
Formate

**Figure S4: Measured volatile fatty acid (VFA) changes during initial hypophosphite dosing. A.** Concentrations of formate (orange), acetate (red), lactate (forest green), propionate (blue), isobutyrate (green), butyrate (yellow), valerate (black), and isovalerate (purple) over time during the initial dosing (0.1mM) of hypophosphite of the treated and untreated designated columns. Error bars represent standard deviation (n=3 per designated condition). **B.** Relative contribution of electrons associated with individual VFAs, expressed as a percentage of the total VFA-measured electrons. Values represent individual columns (1-3 untreated, 4-6 treated). The black line at day five represents the evening that 0.1mM hypophosphite began amendment into the treated columns.

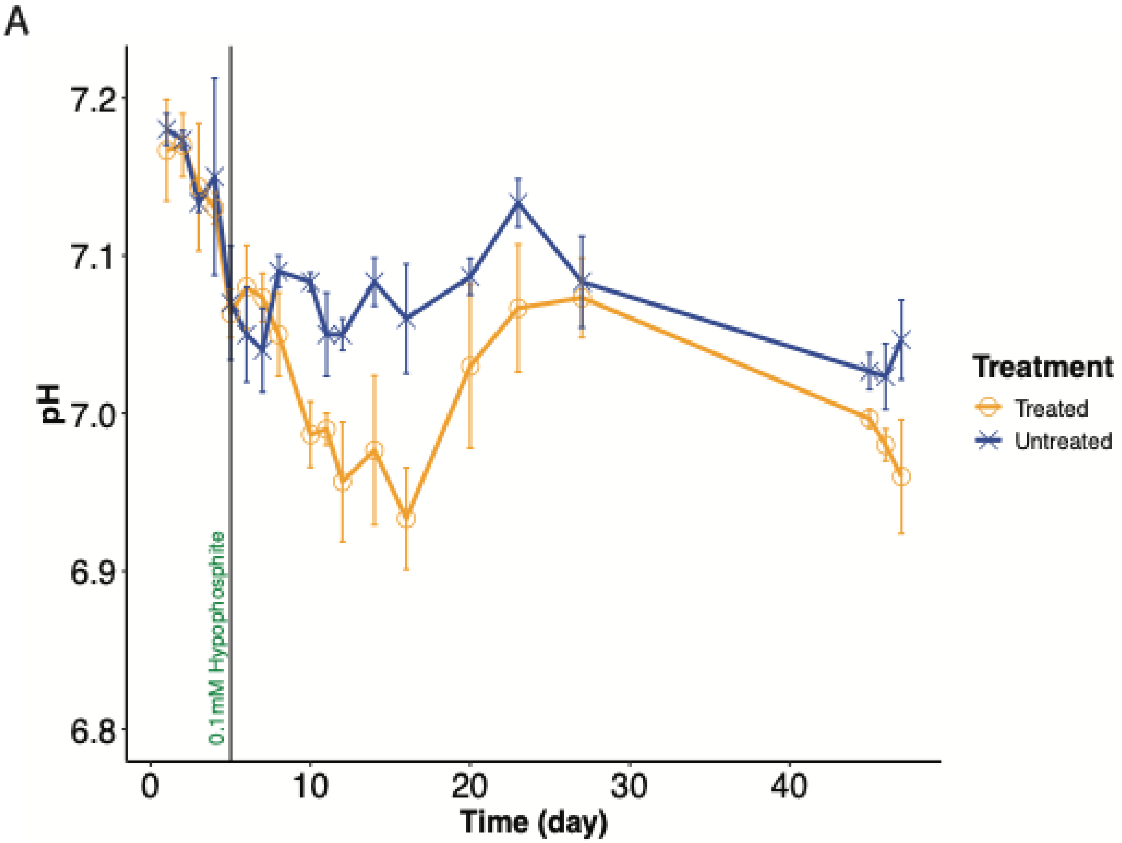
A
pH
7.2
7.1
7.0
6.9
6.8
0.1mM Hypophosphite
0
10
20
30
40
Time (day)
Treatment
Treated
Untreated

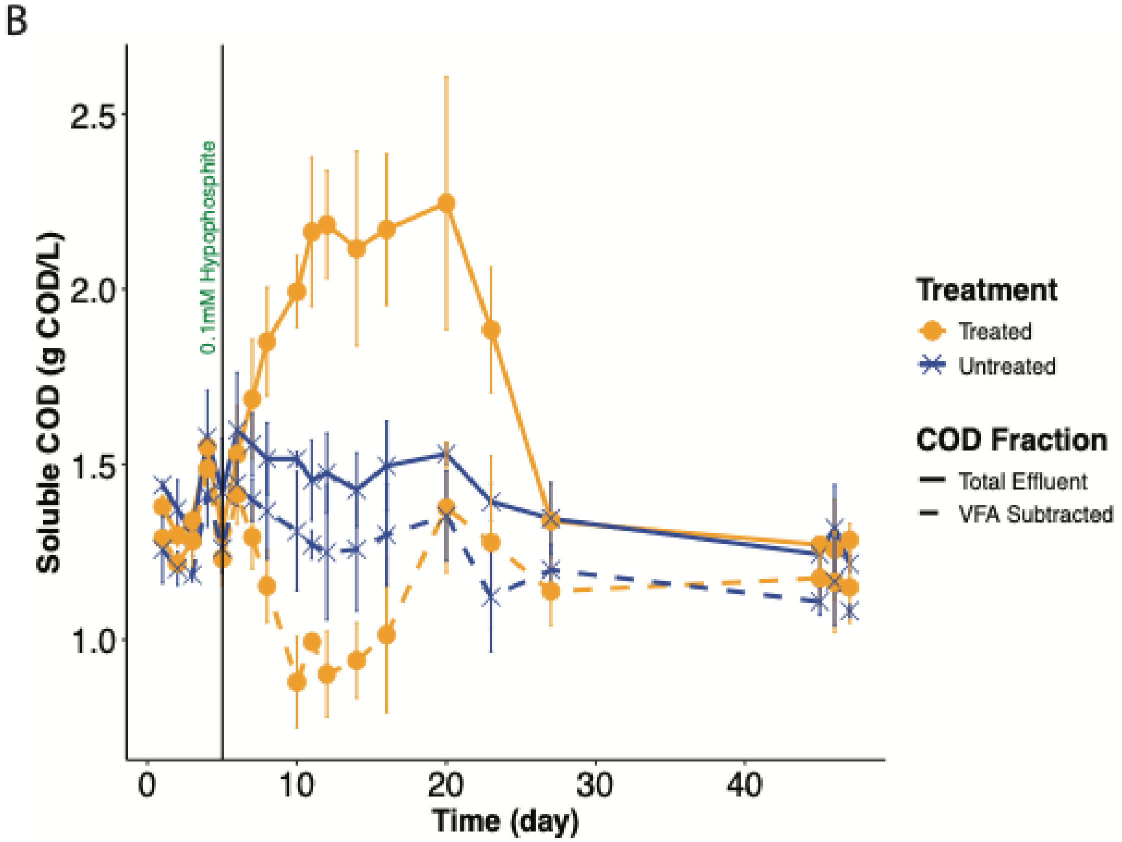
B
Soluble COD (g COD/L)
2.5
2.0
1.5
1.0
0.1mM Hypophosphite
0
10
20
30
40
Time (day)
Treatment
Treated
Untreated
COD Fraction
Total Effluent
VFA Subtracted

**Figure S5: Effects of the initial hypophosphite amendment to the pH and effluent sCOD. A.** pH of treated and untreated columns. **B.** Effluent sCOD (gCOD/L) with and without subtraction of VFA-calculated COD. The (blue x) represents total effluent sCOD and the (orange, solid circles) is effluent sCOD minus VFA COD. Treated columns are shown in orange, solid lines and untreated columns are shown in blue, dashed lines. Error bars represent standard deviation (n=3 per designated condition). The black line at day five represents the evening that 0.1mM hypophosphite began amendment into the treated columns.

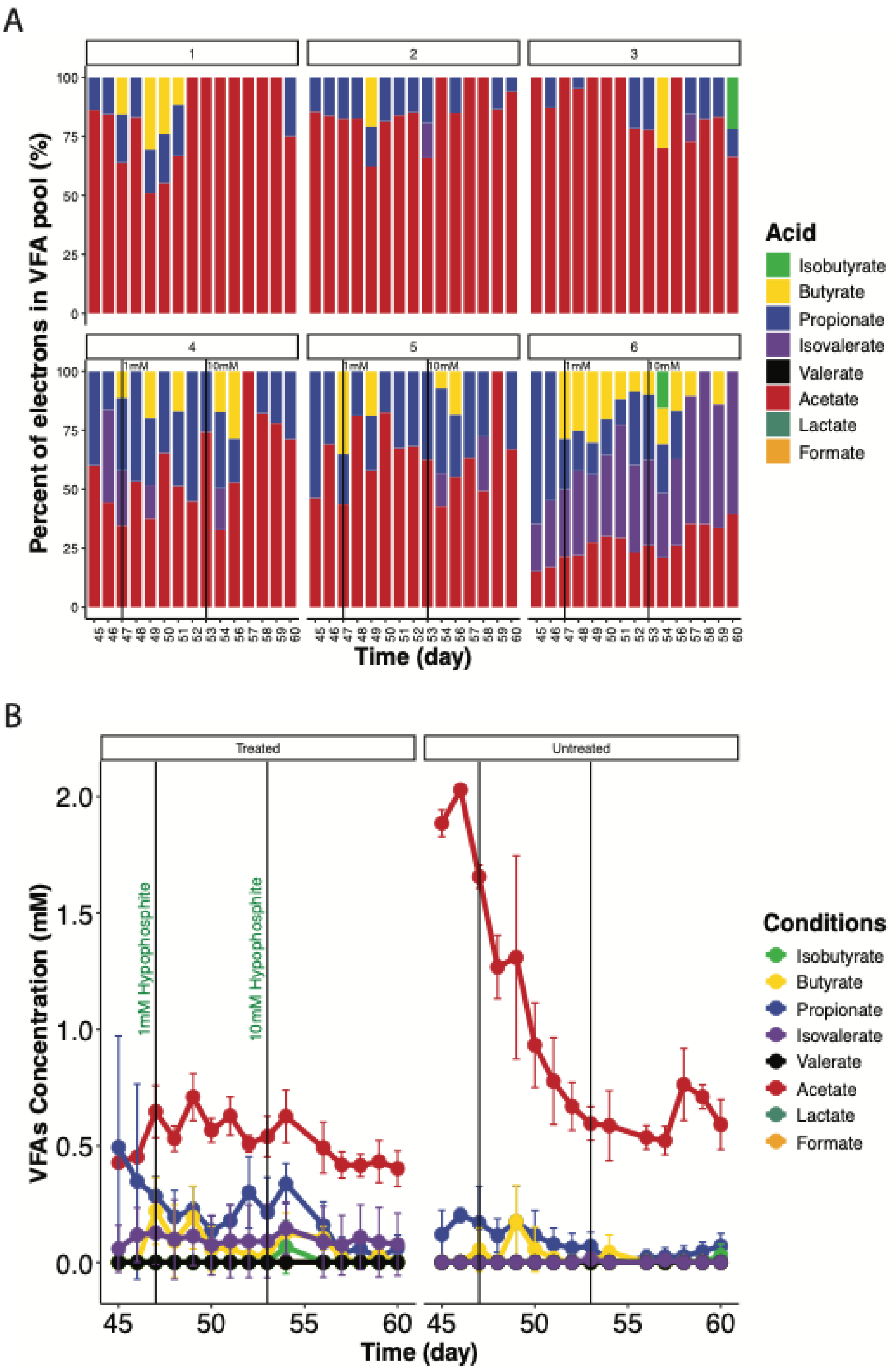
A
1
2
3
4
5
6
1mM
10mM
Percent of electrons in VFA pool (%)
Time (day)
Acid
Isobutyrate
Butyrate
Propionate
Isovalerate
Valerate
Acetate
Lactate
Formate
B
Treated
Untreated
1mM Hypophosphite
10mM Hypophosphite
VFAs Concentration (mM)
Time (day)
Conditions
Isobutyrate
Butyrate
Propionate
Isovalerate
Valerate
Acetate
Lactate
Formate

61

**Figure S6: Measured volatile fatty acid (VFA) changes during 1 and 10mM hypophosphite dosing. A.** Relative contribution of electrons associated with individual VFAs, expressed as a percentage of the total VFA-measured electrons. Values represent individual columns (1-3 untreated, 4-6 treated). **B.** Concentrations of VFAs over time during the second (1mM) and third (10mM) dosing of hypophosphite in the treated. Error bars represent standard deviation (n=3 per condition). The black line at day 47 represents the evening that 1 mM hypophosphite began amendment into the treated columns. The black line at day 53 represents the evening that 10 mM hypophosphite began amendment into the treated columns. Colors for the VFAs are as such: formate (orange), acetate (red), lactate (forest green), propionate (blue), isobutyrate (green), butyrate (yellow), valerate (black), and isovalerate (purple)

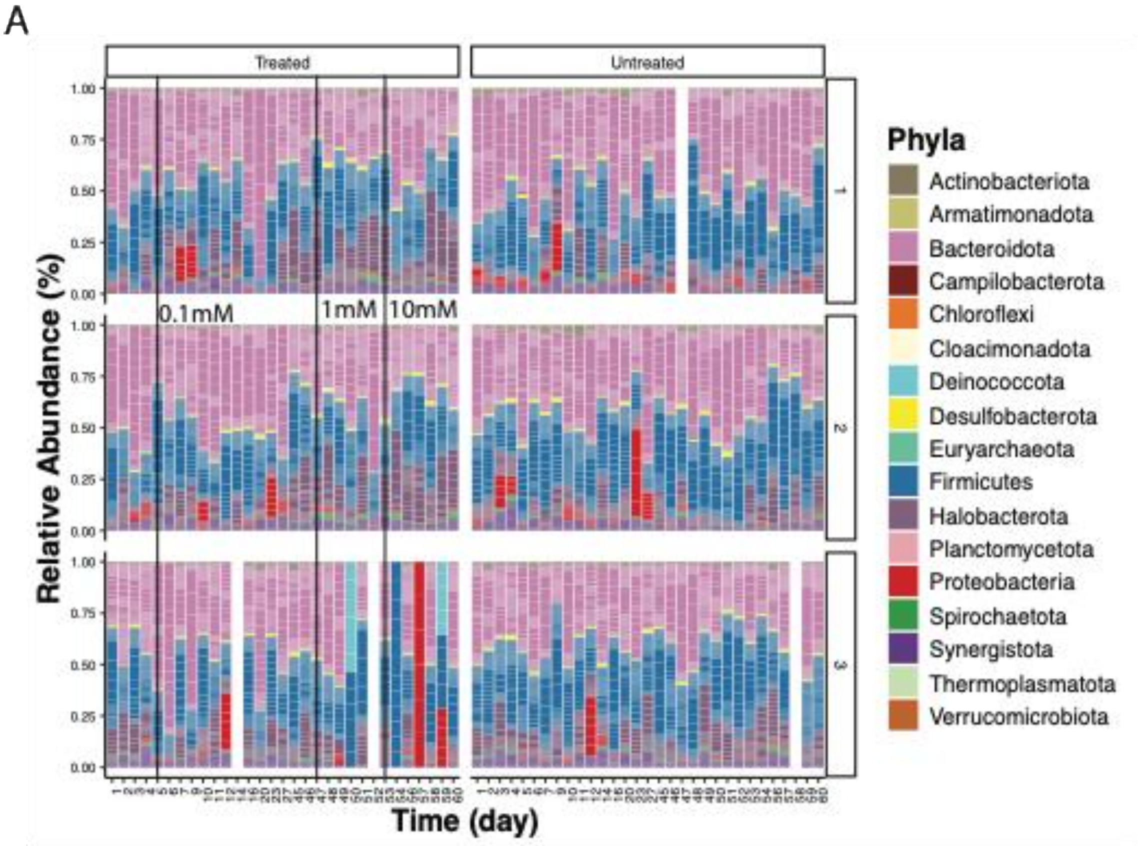


**Figure S7. Temporal changes in UASB microbiome composition in treated and untreated columns during hypophosphite addition. A.** Relative abundance of prokaryotic phyla over the course of the UASB operation in the treated versus untreated columns.

**Supplementary Table Titles**

**Table S1:** Statistical analyses of 16S rDNA amplicon sequencing summarized

**Table S2:** Post-Dunn hoc Statistical analyses on Alpha Diversity Metrics

**Table S3:** Temporal (Phase) comparisons between treatment groups
